# The serine/threonine kinase Back seat driver prevents cell fusion to maintain cell identity

**DOI:** 10.1101/2022.07.22.501168

**Authors:** Shuo Yang, Aaron N. Johnson

## Abstract

Cell fate specification is essential for every major event of embryogenesis, and subsequent cell maturation ensures individual cell types acquire specialized functions. The mechanisms that regulate cell fate specification have been studied exhaustively, and each technological advance in developmental biology ushers in a new era of studies aimed at uncovering the most fundamental processes by which cells acquire unique identities. What is less appreciated is that mechanisms are in place to ensure cell identity is maintained throughout the life of the organism. The body wall musculature in the Drosophila embryo is a well-established model to study cell fate specification, as each hemisegment in the embryo generates and maintains thirty muscles with distinct identities. Here we show the serine/threonine kinase Back seat driver (Bsd), which regulates muscle morphogenesis, also maintains cell identity. Once specified, the thirty body wall muscles fuse with mononucleate muscle precursors that lack a specific identity to form multinucleate striated muscles. Importantly, body wall muscles do not fuse with each other and thereby maintain distinct identities. We show that Bsd prevents inappropriate fusion among the thirty body wall muscles. Thus, the regulation of cell fusion is one mechanism that maintains cell identity.

## Introduction

Cell fate specification is a hallmark of embryonic development, and as development proceeds fated cells mature and acquire specialized identities and functions. Our understanding of cell fate specification was revolutionized by the gene regulatory network (GRN) hypothesis in which extracellular signals regulate the expression of transcription factors that in turn regulate gene expression via *cis*-regulatory modules in the network, establish cell identity, and ultimately activate expression of the structural genes that perform specialized functions (Peter and Davidson, 2016). Cell identity is intimately connected to the overall body plan. Once cells acquire identity and specialized functions, cell identity must be maintained for the organism to operate at the selected capacity. One mechanism that promotes the stability of cell identity involves epigenome regulated changes in the accessibility of chromatin to transcription factors (Balsalobre and Drouin, 2022). However, the genomes of neighboring cells must remain isolated for epigenomics to maintain cell identity.

After gastrulation, the Drosophila mesoderm subdivides into the cardiac, visceral and somatic mesoderm; the somatic mesoderm will give rise to the striated body wall muscles. Thirty distinct body wall muscles develop per hemisegment, and each muscle expresses a distinct set of transcription factors that confers a unique cell identity. The diversification of body wall muscle cell types initiates when Wingless and Decapentaplegic signals establish competence domains in the somatic mesoderm (Carmena et al., 1998). Cells in each domain are competent to respond to subsequent Receptor Tyrosine Kinase signals, which refine the competence domain to a smaller group of equivalent cells. Lateral inhibition within the equivalence group selects a single progenitor to become a founder cell (FC). The remaining cells in the equivalence group will become fusion competent myoblasts (FCMs). Altogether, thirty FCs are specified per hemisegment and each FC acquires a unique identity through the expression of transcription factors known as identity genes (de Joussineau et al., 2012; Sandmann et al., 2006). FCs directionally fuse with FCMs to generate multinucleate myotubes, but FCs will not fuse with each other (Bothe and Baylies, 2016). At the completion of myogenesis, the thirty FCs will have generated thirty body wall muscles with distinct identities, morphologies, and intrasegmental positions. Although the mechanisms that direct FC diversification have been studied in detail, it is unclear how directional myoblast fusion is regulated so that muscle cell identity is maintained.

Here we report that the serine/threonine kinase Back seat driver (Bsd) regulates directional fusion to stabilize muscle cell identity. Live imaging studies revealed multinucleate myotubes derived from distinct FCs fuse with each other in the absence of Bsd activity. We have previously shown that Bsd and the Rho GTPase Tumbleweed (Tum) function in a common pathway to regulate muscle morphogenesis, but Tum does not appear to regulate directional fusion. Instead Bsd regulates Mitogen Activated Protein Kinase (MAPK) activity, which may be a mechanism for limiting non-directional muscle fusion. Interestingly, the transcription factor Jumeau (Jumu) regulates both cell fate specification and chromatin remodeling, and we found Jumu also activates Bsd expression. Our studies suggest that, in addition to epigenetic regulation, Jumu maintains cell identity by preventing cell-cell fusion through a Bsd-dependent mechanism.

## Results and Discussion

Founder cells (FCs) begin to elongate just after specification, and concurrently initiate directional fusion with neighboring fusion competent myoblasts (FCMs) to form syncytial myotubes (Bothe and Baylies, 2016). The nascent myotube leading edges will navigate across the hemisegment and identify muscle attachment sites on tendon cells in the ectoderm. Myotube guidance refers to the combined processes of leading edge navigation and muscle attachment selection, and regulated myotube guidance establishes a stereotypical musculoskeletal pattern in abdominal segments A2-A8 (Figure 1A). We identified Back seat driver (Bsd) in a previous study of myogenesis, and found Bsd is an essential regulator of myotube guidance (Yang et al., 2020).

**Figure 1.**
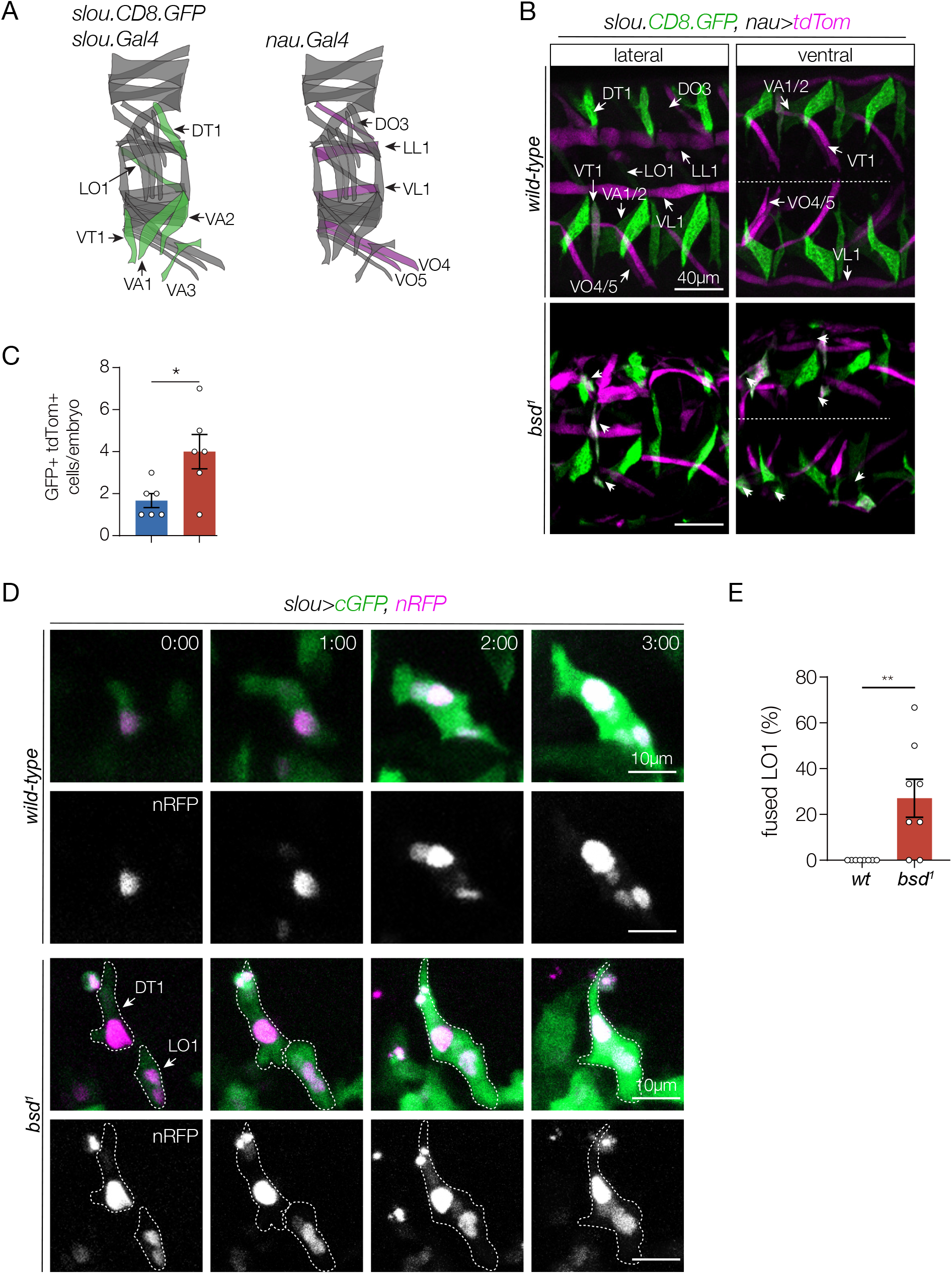
Bsd maintains directional fusion and muscle identity. (A) Diagrams of the stereotypic pattern of body wall muscles in one hemisegment. Each muscle expresses a unique combination of identity genes, including *slouch* (*slou*) and nautilus (*nau*). slou transgenes are active in Doral Transverse 1 (DT1), Longitudinal Oblique (LO1), Ventral Acute 1-3 (VA1-3), and the Ventral Transverse 1 (VT1) muscles (green). A *nau* transgene is active in the Dorsal Oblique 3 (DO3), Longitudinal Lateral 1 (LL1), Ventral Lateral 1 (VL1), and Ventral Oblique 4-5 (VO4-5) muscles (violet). (B) *bsd*^*1*^ myotubes show intermediate identities. Confocal micrographs of live Stage 16 embryos imaged for *slou-CD8-GFP* (green) and *nau>tdTom* (violet). *slou-CD8-GFP* and *nau*.*Gal4>tdTom* are active in largely non-overlapping populations of body wall muscles in wild-type embryos, although VT1 occasionally co-expressed GFP and tdTom. *bsd*^*1*^ embryos had more muscles that co-expressed GFP and tdTom than wild-type embryos (arrowheads). (C) Quantification of GFP/tdTom double positive cells. The number of double positive cells in hemisegments A2-A7 is shown; each data point represents one embryo. (D) *bsd*^*1*^ myotubes undergo non-directional fusion. Live imaging stills of LO1 myotubes in Stage 12-15 embryos that expressed cytoplasmic GFP (green) and nuclear RFP (violet) under the control of *slou*.*Gal4*. Live imaging initiated when GFP fluorescence was detectable in LO1 myotubes (0:00). Control myotubes elongate in a stereotypical fashion and identify attachment sites to acquire an oblique morphology. Notice the number of LO1 myonuclei increases due to directional fusion. *bsd*^*1*^ LO1 myotubes often fused with DT1 myotubes and the LO1-DT1 muscle acquired a transverse morphology. (E) Quantification of non-directional myotube fusion from live imaging. The number of LO1 fusion events with other *slou>GFP* myotubes in hemisegments A2-A7 is shown; each data point represents one embryo. Significance in (C,E) was determined by unpaired students t-test. Error bars represent SEM. (*) p<0.05, (**) p<0.01. (#:##) hr:min.

Each FC expresses a unique set of transcription factors known as identity genes. Enhancers from the identity gene *slouch* (*slou*) were used to generate a transgenic reporter gene (*slou-CD8-GFP*), which is active in six FCs that give rise to the Dorsal Transverse 1 (DT1), the Longitudinal Oblique 1 (LO1), Ventral Acute 1-3 (VA1-3), and the Ventral Transverse 1 (VT1) muscles (Figure 1A)(Schnorrer et al., 2007). A transgenic *Gal4* line constructed from similar *slou* enhancers (*slou*.*Gal4*) recapitulates the activity of the *slou-CD8-GFP* reporter (Yang et al., 2020). Enhancers from the identity gene *nautilus* (*nau*) comprise a third transgenic line (*nau*.*Gal4*), which is active in the Dorsal Oblique 3 (DO3), Longitudinal Lateral 1 (LL1), Ventral Lateral 1 (VL1), and Ventral Oblique 4-5 (VO4-5; Fig. 1A)(McAdow et al., 2022). *slou-CD8-GFP* and *nau*.*Gal4* are active in non-overlapping populations of body wall muscles, and in wild-type embryos *slou-CD8-GFP* and *nau*.*Gal4, UAS*.*tdTomato* (*nau>tdTom*) were rarely co-expressed in body wall muscles (Fig. 1B,C). Surprisingly the number of muscles that co-expressed *slou-CD8-GFP* and *nau>tdTom* in *bsd*^*1*^embryos was significantly greater than controls (Fig. 1B,C). These data suggest *bsd*^*1*^ muscles adopt abnormal identities.

FCs are correctly specified in *bsd*^*1*^ embryos (Yang et al., 2022), so we used live imaging to understand how Bsd might regulate muscle identity. Nascent myotubes extend bipolar projections to connect with muscle attachment sites that link muscles to the exoskeleton (Fig. 1D, Supplementary Movie 1). During elongation, LO1 myotubes that expressed *slou>GFP* fused with FCMs but not with other myotubes that expressed *slou>GFP* (Fig. 1E). In contrast, 28% of LO1 myotubes in *bsd*^*1*^ embryos fused with neighboring *slou>GFP* myotubes (Fig 1D,E), arguing Bsd prevents non-directional fusion between myotubes to maintain cell identity.

Bsd directly activates Polo kinase, and active Polo in turn regulates many downstream effectors, including the GTPase activating protein Tumbleweed (Tum) and the kinesin microtubule motor protein Pavarotti (Pav) (D’Avino et al., 2006; Ebrahimi et al., 2010; Gregory et al., 2008; Somers and Saint, 2003). Polo, Tum, and Pav act in a common myogenic pathway to direct myotube guidance (Guerin and Kramer, 2009; Yang et al., 2022), so we asked if Bsd acts through the Polo/Tum/Pav cytoskeletal regulatory module to maintain direction fusion. *tum*^*DH15*^ produced the strongest muscle phenotypes among the *polo, pav*, and *tum* alleles we tested (Yang et al., 2022) and, while live imaging showed *tum*^*DH15*^ embryos have highly penetrant myotube guidance defects, we did not observe any instances of LO1 myotubes fusing with neighboring *slou>GFP* myotubes (Fig 2A,B; Supplementary Movie 2). These live imaging studies argue Bsd acts through a Polo/Tum/Pav independent pathway to regulate cell fusion.

**Figure 2.**
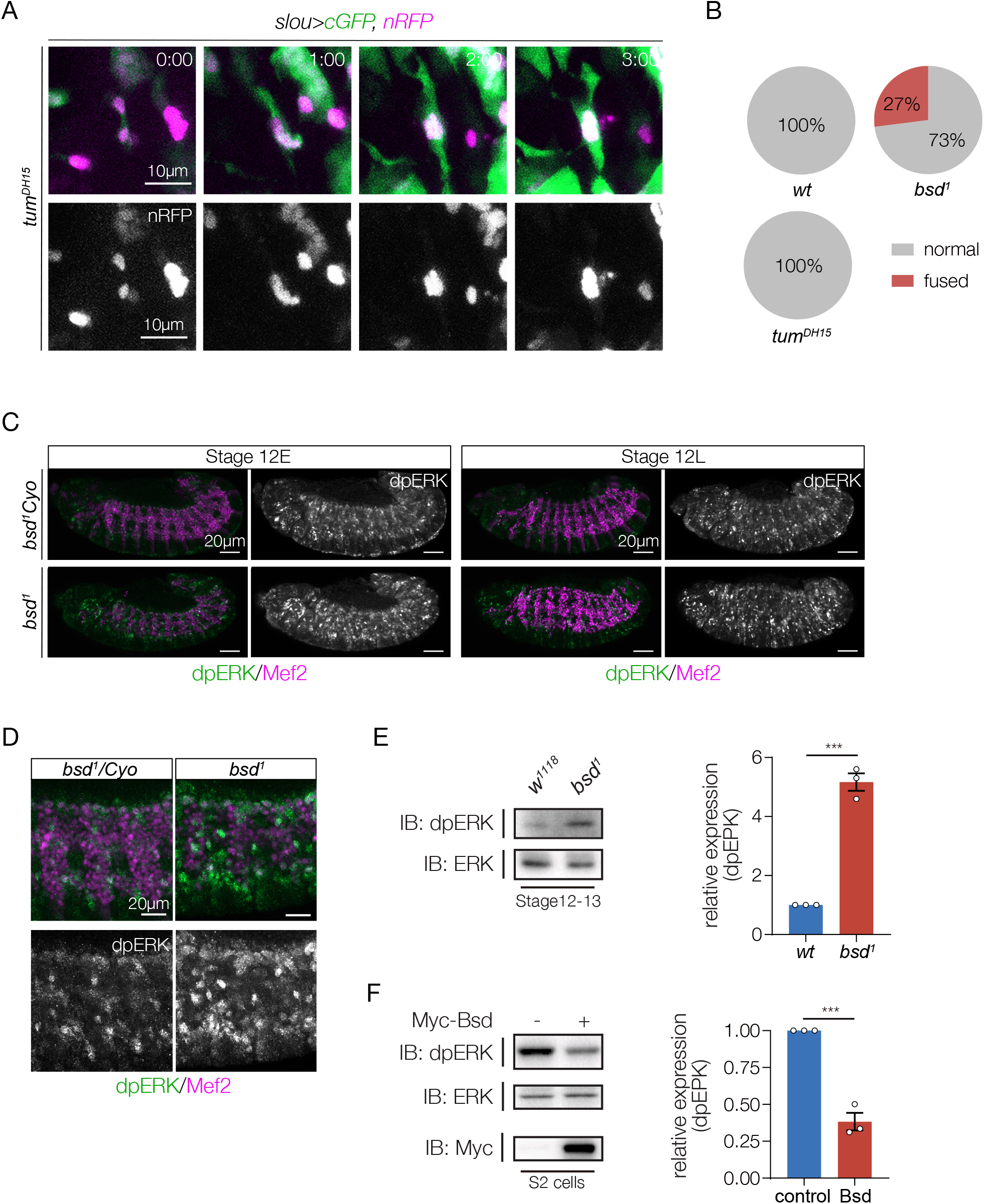
Bsd regulates ERK activity. (A) Tum does not regulate directional fusion. Live imaging stills of LO1 myotubes in Stage 12-15 embryos as described in Fig. 1. *tum*^*DH15*^ LO1 myotubes showed severe guidance defects, but did not fuse with other *slou>GFP* myotubes. (B) Quantification of non-directional myotube fusion from live imaging. n≥6 embryos per genotype. (C,D) Bsd represses ERK phosphorylation. Stage 12 embryos labeled for dpERK (green) and Mef2 (violet). dpERK identifies cells with activated ERK and Mef2 is expressed in all myogenic cells. *bsd*^*1*^ embryos showed more dpERK in the mesoderm than controls. High magnification confocal micrographs of the embryos in (C) are shown in (D). (St12E) Stage 12 early, (St12L) Stage 12 late. (E) Stage 12 embryo lysates immunoblotted with dpERK. *bsd*^*1*^ embryo lysates showed significantly more phosphorylated ERK than controls. (F) Cells transfected with Bsd showed significantly less phosphorylated ERK than controls. Lysates from S2 cells transfected with Bsd.Myc and control plasmids were immunoblotted with dpERK and Myc to detect transgenic proteins. Data points in the dpERK quantifications (E,F) represent one biological replicate. Significance was determined by unpaired students t-test. Error bars represent SEM. (***) p<0.001.

In Drosophila visceral muscles, Anaplastic lymphoma kinase (Alk) activates extracellular signal-regulated kinase (ERK) to induce the expression of the cell adhesion molecule Kirre, which is essential for myoblast fusion (Englund et al., 2003; Krauss, 2010). In contrast, the Bsd orthologue Vaccinia-related kinase 3 (VRK3) inactivates ERK by regulating the ERK phosphatase Vaccinia H1-related (Kang and Kim, 2006, 2008). One possibility is that Bsd inactivates ERK in Drosophila to suppress Kirre expression, and in turn regulate cell fusion. We assayed phosphorylated ERK (dpERK) *in vivo* and *in vitro*, and found *bsd*^*1*^ embryos had significantly more dpERK than control embryos (Fig. 2C-E), and S2 cells transfected with Bsd had significantly less phosphorylated ERK than control transfected cells (Fig. 2F). These results are consistent with the hypothesis that Bsd inactivates ERK to regulate cell fusion. Although a Kirre antibody is no longer available, our studies suggest that ERK-regulated pathways control Kirre expression in body wall muscles to direct myoblast fusion.

We next asked what pathways might be regulating Bsd to ensure muscle cell identity is correctly maintained. Using mass spectrometry, we identified a number of proteins that physically interact with Bsd, including Polo, but none of the candidates are known regulators of myogenesis (Yang et al., 2022). Regulatory kinases are differentially expressed during organogenesis (Yang et al., 2022), so we used the DNAMAN prediction tool to identify transcription factor binding sites near the *bsd* locus. *bsd* transcription is controlled by two core promoters, and each promoter is less than 400bp from a consensus binding site for the transcription factor Jumeau (Jumu; Fig 3A). Chromatin immunoprecipation and massively parallel DNA sequencing (ChIP-seq) for Jumu in 0-24hr embryos was performed by the modENCODE project (ENCSR946VDB)(Luo et al., 2020), which identified a Jumu binding region that overlaps with the *bsd* core promoters (Fig. 3A). To understand if Jumu regulates *bsd* expression, we optimized a quantitative real time PCR (qRT-PCR) strategy by testing the expression of a known Jumu target gene, *heartless* (*htl*), in embryos homozygous for a strong hypomorphic allele of *jumu*^*2*.*12*^ (Fig. 3B)(Ahmad et al., 2016). qRT-PCR showed that both *htl* and *bsd* expression was significantly reduced in *jumu*^*2*.*12*^ embryos compared to controls (Fig. 3B). In addition, *jumu*^*2*.*12*^ embryos showed muscle morphogenesis defects similar to *bsd*^*1*^ embryos (Fig. 3C), and *slou>GFP* myotubes were short and incorrectly targeted in *jumu*^*2*.*12*^ embryos (Fig. 3D). Our studies are consistent with a model in which Jumu directly activates *bsd* expression to regulate myotube guidance and maintain muscle cell identity. Jumu also acts upstream of Polo through an unknown mechanism to regulate cardiac muscle specification (Ahmad et al., 2012). One exciting possibility is that Jumu transcriptionally activates *bsd* to regulate Polo activity.

**Figure 3.**
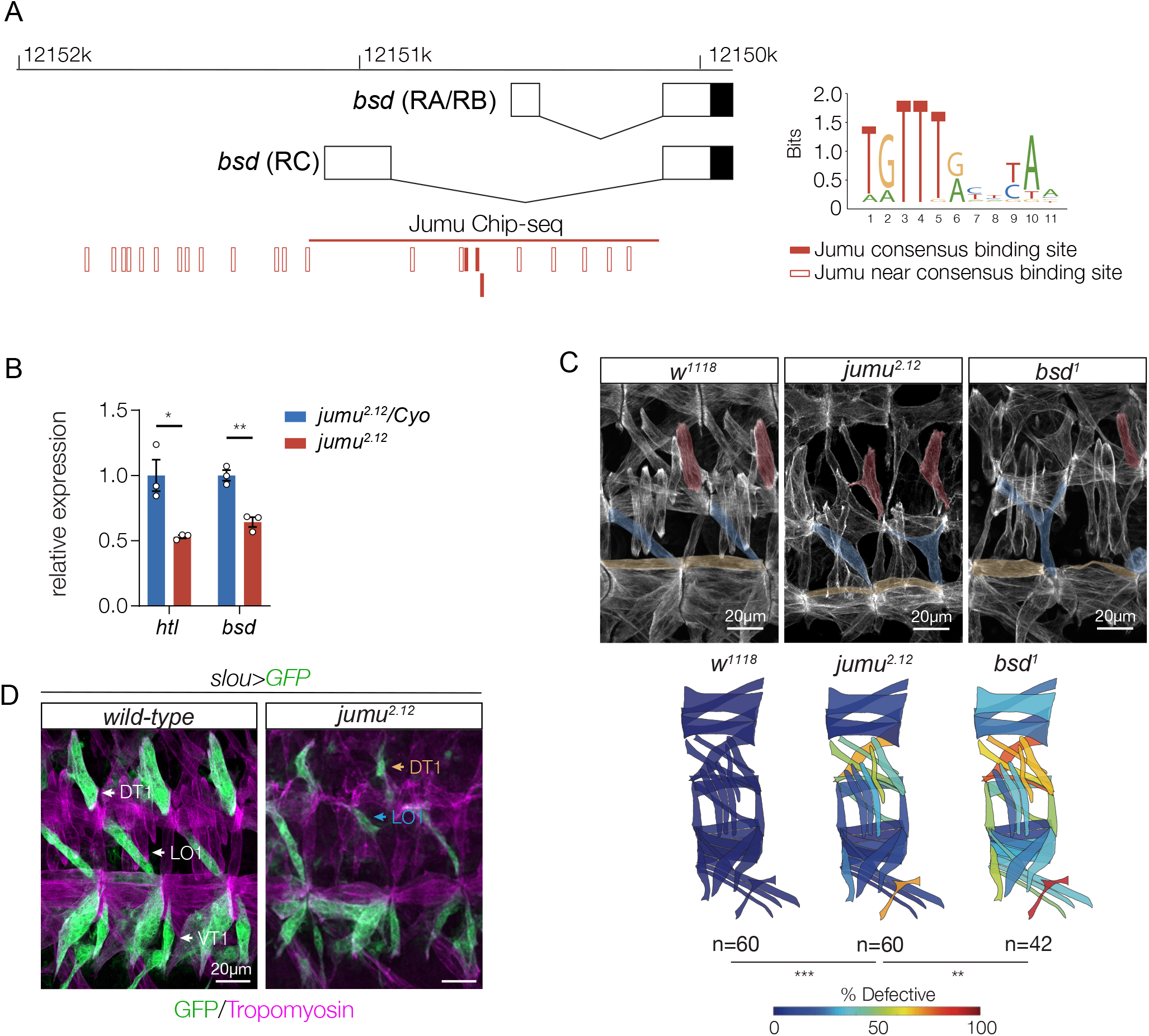
Jumu activates *bsd* expression. (A) Diagram of the *bsd* core promoter region. Open boxes represent untranslated regions; shaded boxes show the beginning of the open reading frame. Molecular coordinates refer to chromosome 2R. The consensus Jumu binding site was reported in (Hao and Jin, 2017). The ChIP-seq binding region was reported by modENCODE project (ENCSR946VDB). (B) Quantitative real time PCR analysis of *htl* and *bsd* mRNA showed each transcript was reduced in Stage 12 *jumu* embryo lysates (n=3 biological replicates). (C) Muscle phenotypes in *jumu*^*2*.*12*^ and *bsd*^*1*^ embryos. Stage 16 embryos labeled with Tropomyosin. DT1, LO1, and VL1 muscles are pseudocolored red, blue, and yellow. Muscle morphology defects was comparable between *jumu*^*2*.*12*^ and *bsd*^*1*^ embryos. Quantification of muscle phenotypes. Muscles were scored in hemisegments A2-A7 of St16 embryos. Abnormal phenotypes (missing muscles, muscles with attachment site defects, and muscles that failed to elongate) were scored “defective”. The frequencies of muscle defects are shown as a heat map on the stereotypic muscle pattern in one embryonic hemisegment. (n) number of hemisegments scored. (D) *jumu*^*2*.*12*^ DT1, LO1, and VT1 muscle phenotypes. Stage 16 embryos labeled for *slou>GFP* (green) and Tropomyosin (violet). *jumu*^*2*.*12*^ muscles had attachment site defects (orange and blue arrows). Significance was determined by unpaired students t-test (B) or Fisher’s exact test (C). Error bars represent SEM. (*) p<0.05, (**) p<0.01, (***) p<0.001.

The specification of thirty distinct founder cells (FCs) per hemisegment is accomplished through morphogen-induced specification of equivalence groups, lateral inhibition within equivalence groups to establish individual progenitors, and combinatorial expression of distinct transcription factors to generate diverse cell identities. The final musculoskeletal pattern however is established when FCs undergo a morphological transformation into myotubes, and respond to navigational cues transduced by the transmembrane receptors Heartless, Kon-tiki, and Robo to elongate and identify muscle attachment sites at the segment boundary (Kramer et al., 2001; Schnorrer et al., 2007; Yang et al., 2020). The developmental mechanisms that direct FC cell fate specification and morphogenesis are commonly used strategies that regulate organogenesis across Metazoa, but FCs also employ a muscle-specific developmental program, directional fusion, to generate multinucleate muscle cells with distinct identities. FCs and myotubes only fuse with mononucleate fusion competent myoblasts (FCMs), but do not fuse with each other (Fig. 1D)(Bothe and Baylies, 2016).

Directional fusion persists during myotube guidance through a Bsd-dependent mechanism. Myotubes seeded from distinct FCs do not fuse with each other, but in *bsd* mutant embryos the loss of directional fusion produced muscles with intermediate identities (Fig. 1D). Directional fusion does not appear to be regulated by the position of the myotube in the segment. *tum* and *htl* myotube leading edges navigate to incorrect positions, but do not fuse with other myotubes (Fig. 2A)(Yang et al., 2020). These observations suggest Bsd prevents myotube fusion by blocking the activity or function of pro-fusion pathways in maturing myotubes. The nephrin-like transmembrane proteins Kirre and Sticks and stones (Sns) are differentially expressed in muscle precursors: Kirre is expressed in FCs and Sns is expressed FCMs (Bour et al., 2000; Ruiz-Gómez et al., 2000). Sns is a ligand of Kirre, and heterophilic interactions between Kirre and Sns are thought to confer FC-FCM cell recognition, and provide the adhesive forces necessary for fusion (Krauss, 2010). Homophilic interactions between Kirre expressing cells in culture promotes cell-cell adhesion (Menon et al., 2005), suggesting enhanced expression or activation of Kirre in myotubes could drive non-directional fusion. *Kirre* transcription is induced in visceral muscle cells by ERK signaling (Englund et al., 2003), and we show Bsd negatively regulates ERK activation (Fig. 2C-F). It is possible that Bsd suppresses Kirre activity to maintain directional fusion and preserve muscle cell identity. However, spatial misexpression of Kirre in the somatic mesoderm alone is not sufficient to induce obvious non-directional cell fusion (Menon et al., 2005), suggesting Bsd regulates multiple cell fusion proteins.

The expression of regulatory kinases is spatially and temporally dynamic, and approximately 50% of Drosophila kinases show tissue-specific expression during the late stages of embryogenesis (Yang et al., 2022). Bsd expression follows this pattern, and is enriched in the somatic mesoderm during myotube guidance (Yang et al., 2022). Htl is a receptor tyrosine kinase that transduces FGF signals, and Htl is broadly expressed in the mesoderm after gastrulation (Fisher et al., 2012). However, during organogenesis, *htl* expression is restricted to a subset of progenitors that will give rise to cardiac, somatic, and visceral muscle (Yang et al., 2020). We confirmed that Jumu regulates *htl* expression in Stage 12 embryos, and show Jumu also regulates *bsd* expression (Fig. 3A,B). Jumu is expressed in FCs and, although repressors have been identified that block *jumu* expression in FCMs, the mechanisms that directly activate *jumu* expression in the mesoderm are unknown (Ahmad et al., 2012; Ciglar et al., 2014). Nonetheless, our study shows that Jumu dictates the expression of at least two regulatory kinases, Bsd and Htl, which argues only a small number of transcription factors may be needed to establish the dynamic spatiotemporal expression of regulatory kinases that control cell morphogenesis. Since zebrafish regulatory kinases also show dynamic expression patterns (Yang et al., 2022), it will be important to understand how cell identity and the expression of morphogenetic transcription factors like Jumu are coordinated to direct organogenesis across Metazoa.

## Acknowledgements

We thank Shaad Amhad (Indiana State University) for sending *jumu* flies, and Frank Schnorrer (IBDM, Marseille, France) for providing *slou-mCD8-GFP* flies. We also thank the larger Drosophila community for stocks and reagents, and Kevin White’s laboratory for performing the Jumu ChIP-seq. ANJ was supported by NIH R01AR070299.

## Author Contributions

Conceptualization: S.Y. and A.N.J., Methodology: S.Y. and A.N.J., Formal analysis: S.Y. and A.N.J., Investigation: S.Y., Resources: A.N.J.; Data curation: S.Y., Writing original draft: S.Y. and A.N.J.

## Methods

### *Drosophila* genetics

The following stocks were obtained from the Bloomington Stock Center: *tum*^*DH15*^, *P{GMR40D04-GAL4}attP2* (*slou*.*Gal4*), and *P{GMR57C12-GAL4}attP2* (*nau*.*Gal4*). The other stocks used in this study were *bsd*^*1*^ (Yang et al., 2022), *jumu*^*2*.*12*^ (Ahmad et al., 2012), and *P{slou-mCD8-GFP} (Schnorrer et al*., *2007). Cyo, P{Gal4-Twi}, P{2X-UAS*.*eGFP}; Cyo, P{wg*.*lacZ}; TM3, P{Gal4-Twi}, P{2X-UAS*.*eGFP}; and TM3, P{ftz*.*lacZ}* balancers were used to genotype embryos.

### Immunohistochemistry

Antibodies used were α -Mef2 (1:1000, gift from R. Cripps), α -Tropomyosin (1:600, Abcam, MAC141), α -GFP (1:600, Torrey Pines Biolabs, TP-401), and α -dpERK (1:300, Cell Signaling Technologies, 4377). Embryo antibody staining was performed as described (Johnson et al., 2013); HRP-conjugated secondary antibodies in conjunction with the TSA system (Molecular Probes) were used to detect primary antibodies.

### Imaging and image quantification

Embryos were imaged with a Zeiss LSM800 confocal microscope. For time-lapse imaging, dechorionated St12 embryos were mounted in halocarbon oil and scanned at 6min intervals. Control and mutant embryos were prepared and imaged in parallel where possible, and imaging parameters were maintained between genotypes. Fluorescent intensity and cell morphology measurements were made with ImageJ software.

### Phenotypic scoring, analysis, and visualization

Each embryonic hemisegment has 30 distinct muscles with a fixed pattern as shown in Figure 1A. Muscle phenotypes were analyzed in hemisegments A2-A7, in a minimum of nine embryos per genotype. A Tropomyosin antibody was used to visualize all body wall muscles, and the percent defective was calculated for each of the 30 muscles in minimum of 42 hemisegments from seven different embryos. <u>% Defective</u> = # of abnormal muscles/hemisegments scored. To visualize affected muscles, percent defective was converted to a schematic heat map on the body wall muscle pattern.

### Immunoprecipitation and Western blotting

For *Drosophila* proteins, S2 cells (8×10^6^) were transfected with 1.5µg of pAMW.Bsd or pAMW plasmids in 6-well plates. Cells were cultured for 24h collected, washed twice with PBS, lysed with 600µl IP buffer (20 mM Hepes, pH=7.4, 150 nM NaCl, 1% NP40, 1.5 mM MgCl_2_, 2 mM EGTA, 10 mM NaF, 1 mM Na_3_VO_4_, 1X proteinase inhibitor), incubated on ice for 30 min, centrifuged at 12000Xg for 15min. The supernatant was collected, and Western blots were performed by standard method using precast gels (#456-1096, BioRad), and imaged with the ChemiDoc XRS+ system (BioRad). Antibodies used for Western blots were α -dpERK (1:1000, Cell Signaling Technologies, 4377), α -ERK (1:1000, Enzo Life Sciences, ADI-KAP-MA001), and α -Myc (1:1000, Sigma, PLA0001).

### Quantitative real time PCR

Total RNA was extracted with RNeasy mini kit (74104, Qiagen), and quantified (Nanodrop 2000). cDNA was prepared by reverse transcription with M-MLV Reverse Transcriptase (28025013, Thermo) with 2000ng RNA. PowerUp Sybr Green Master Mix (A25742, Thermo) and ABI StepOne system (Applied Biosystems) were used for quantitative RT-PCR. Quantification was normalized to *GAPDH* or *RpL32*. Primers used: Htl-F-5’-ACCAAATTGCCAGAGGAATG-3’ Htl-R-5’-GGTAGCCTGCCATTTGTGTT-3’ Bsd-F-5’-TCAACGCTAAGCACTCCGTT-3’ Bsd-R-5’-CGCCTCTGCTCCATGTCTAG-3’ Rp32-F-5’-ATGCTAAGCTGTCGCACAAATG-3’ Rp32-R-5’-GTTCGATCCGATACCGATGT-3’

### Bioinformatic and statistical analysis

DNAMAN (Lynnon Biosoft, Ver. 10) was used to identify consensus transcription factor binding sites. Statistical analyses were performed with GraphPad Prism 9 software, and significance was determined with the unpaired student’s t-test, and two-sided Fisher’s exact test. Sample sizes are indicated in the figure legends. All individuals were included in data analysis.

## Movie Captions

**Supplementary Movie 1. Bsd prevents non-directional fusion**. Live imaging of LO1 myotubes from Stage 12 *slou>eGFP,nRFP* embryos. *slou> eGFP* (green) myotubes fused in *bsd*^*1*^ embryos but not controls. The number of nRFP positive myonuclei (violet) increased due to directional fusion between myotubes and fusion competent myoblasts.

**Supplementary Movie 2. Tum does not regulate myotube fusion**. Live imaging of LO1 myotubes from Stage 12 *tum*^*DH15*^ *slou>eGFP,nRFP* embryos. *slou> eGFP* (green) myotubes did not fuse in *tum*^*DH15*^ embryos. Interestingly, the myonuclei (violet) remained clustered in mutant myotubes.

